# Zika virus infection during pregnancy protects against secondary infection in the absence of CD8+ cells

**DOI:** 10.1101/2020.05.08.082610

**Authors:** Blake Schouest, Margaret H. Gilbert, Rudolf P Bohm, Faith Schiro, Pyone P. Aye, Antonito T Panganiban, Diogo M. Magnani, Nicholas J Maness

**Affiliations:** Tulane National Primate Research Center, Tulane University, Covington, Louisiana, USA; Biomedical Sciences Training Program, Tulane University School of Medicine, New Orleans, Louisiana, USA; Department of Microbiology and Immunology, Tulane University School of Medicine, New Orleans, Louisiana, USA; Department of Medicine, University of Massachusetts, Boston, Massachusetts, USA

**Keywords:** Zika virus, pregnancy, nonhuman primates, macaques, CD8+ T cells

## Abstract

While T cell immunity is an important component of the immune response to Zika virus (ZIKV) infection generally, the efficacy of these responses during pregnancy remains unknown. Here, we tested the capacity of CD8 lymphocytes to protect from secondary challenge in four macaques, two of which were depleted of CD8+ cells prior to rechallenge with a heterologous ZIKV isolate. The initial challenge during pregnancy produced transcriptional signatures suggesting complex patterns of immune modulation, but all animals efficiently controlled the rechallenge virus, implying that the primary infection conferred adequate protection. The secondary challenge promoted humoral responses and activation of innate and adaptive immune cells, suggesting a brief period of infection prior to clearance. These data confirm that ZIKV infection during pregnancy induces sufficient immunity to protect from a secondary challenge and suggest that this protection is not solely dependent on CD8 T cells but entails multiple arms of the immune system.

## Introduction

ZIKV was first isolated nearly seventy years prior to the Brazilian outbreak of 2015 (Dick et al., 1952; Zanluca et al., 2015), but the recent epidemic became associated with vertical transmission dynamics and congenital syndromes that were unprecedented for ZIKV or any other flavivirus (Plourde and Bloch, 2016). Although infrequent neurological manifestations, including Guillain-Barre syndrome, meningitis, and meningoencephalitis, became linked to infection in adults (Araujo et al., 2016; Avelino-Silva and Martin, 2016; Brasil et al., 2016; Ellul et al., 2016), the most severe neurological consequences were documented in infants born to mothers infected during pregnancy (Barton and Salvadori, 2016; Melo et al., 2016). Referred to as congenital Zika syndrome (CZS) (Moore et al., 2017), this collection of manifestations has provided the greatest justification to develop prophylactic and therapeutic countermeasures against the virus. Several murine and NHP models have been developed to understand mechanisms of maternal-to-fetal transmission and to develop and test antiviral therapies and vaccines (Aliota et al., 2016; Dudley et al., 2016; Hirsch et al., 2017; Koide et al., 2016; Magnani et al., 2018; Morrison and Diamond, 2017; Nguyen et al., 2017; O’Connor et al., 2018; Osuna et al., 2016), but NHPs may provide a superior model to study vertical transmission and congenital hazards due to the similarities in placental structure and gestational development to humans (Morrison and Diamond, 2017).

Experimental ZIKV vaccine efforts to date have been successful, with a number of candidate vaccines having advanced to clinical trials, but an often underappreciated consideration in vaccine design is whether protective responses can be attained in the context of pregnancy. Complex interactions between sex hormones and the immune system make pregnant women more susceptible to a host of infections (Kourtis et al., 2014), so an important question in ZIKV vaccine design is whether immunity induced during pregnancy is sufficient to prevent subsequent infections and if this protection extends to infants born to women infected during pregnancy.

A recent study showed that NHPs infected during pregnancy establish long-term immune responses that are sufficient to protect against secondary challenge (Moreno et al., 2019), a finding that is also true in non-pregnant macaques (Aliota et al., 2016). Similar to other flaviviruses, ZIKV infection results in rapid neutralizing antibody titers (Coffey et al., 2017), suggesting that humoral immunity may be the most important correlate of protection. However, ZIKV-specific T cell responses have been described in mice (Elong Ngono et al., 2017; Huang et al., 2017; Pardy et al., 2017), macaques (Dudley et al., 2016), and humans (Grifoni et al., 2018; Grifoni et al., 2017; Ricciardi et al., 2017; Xu et al., 2016), so cell-mediated immunity might also have role in protection from secondary infection. Here, we used the rhesus macaque model to address whether ZIKV infection during pregnancy induces sufficient immunity to protect from rechallenge, and we also asked whether CD8 lymphocytes are an important component of this protection.

## Materials and Methods

### CD8 depletion

Approximately nine months after an initial challenge during the 3^rd^ trimester of pregnancy, as described previously (Magnani et al., 2018), two of four dams were depleted of CD8α+ lymphocytes (primarily NK cells and CD8+ T cells) using the anti-CD8α MT807R1 antibody (Nonhuman Primate Reagent Resource, RRID:AB_2716320) with a standard four-dose regimen over 10 days. CD8+ cell counts in blood were monitored by FACS analysis, and animals were screened for adverse events after each administration and none were observed.

### Viral challenge and viral load quantification

Primary ZIKV inoculations of the primary ZIKV isolate Rio-U1 were described previously (Magnani et al., 2018). The challenge virus was isolated in 2015 from the urine of a pregnant woman in Rio de Janeiro (Bonaldo et al., 2016). Briefly, four Indian rhesus macaques were initially challenged with ZIKV Rio-U1 at 10^4^ plaque forming units (PFU) during the 3^rd^ trimester of pregnancy, resulting in serum viremia that peaked at 3 days post infection (dpi) with 5-6 logs of viral RNA/ml in plasma that cleared between 14 and 28 dpi. One animal, M09, had detectable virus in amniotic fluid just prior to full term fetal harvest, between 35 and 40 dpi. Two infants were sacrificed for tissue harvest, but no evidence of *in utero* infection was present. The other two infants were kept alive for a viral challenge and behavioral observation, as described previously (Maness et al., 2019).

Secondary inoculations of the heterologous Puerto Rican isolate PRVABC-59 were carried out at the same dose (10^4^ PFU), and route (subcutaneous) as the primary challenge. Blood and cerebrospinal fluid (CSF) were drawn on days 0, 3, 5, 7, 14, and 28 post challenge. Viral RNA was isolated from serum and CSF using the Roche High Pure Viral RNA Kit followed by quantification as described previously (Magnani et al., 2018). Animals were euthanized 28 dpi (n=2) or 30 dpi (n=2) after secondary challenge.

### RNA-sequencing and analysis

Total RNA was extracted from PBMC pellets at the indicated timepoints using the Zymo Quick-RNA Miniprep kit. RNA was purified using the Zymo RNA clean & concentrator-25 kit and quantitated using the Qubit RNA BR assay kit (Thermo Fisher). A beta release of the Collibri 3’ mRNA Library Prep Kit (Invitrogen) was used to prepare libraries, and sequencing was carried out at the Tulane NextGen sequencing core using an Illumina NextSeq instrument with 150 cycles.

Sequencing data were aligned and mapped to the rhesus macaque genome (Mmul_10 assembly) using STAR (Dobin et al., 2013) with default settings in gene quantification mode. Differentially expressed genes (DEGs) were calculated using DESeq2 (Love et al., 2014), and pathway analysis was carried out using gene set variation analysis (GSVA) (Hänzelmann et al., 2013), gene set enrichment analysis (GSEA) (Subramanian et al., 2005), ReactomePA (Yu and He, 2016), and Ingenuity Pathway Analysis (IPA) (Qiagen). For GSVA analysis, gene sets in the Reactome databank were used, and pairwise comparisons among conditions were carried out using limma (Ritchie et al., 2015). Gene sets were considered significantly differentially enriched at p<0.05. For GSEA analysis, a false discovery rate (FDR) below 25% was used to identify gene sets in the Hallmarks collection that were significantly enriched at 3 or 7 dpi relative to pre-infection. For these analyses, a gene set permutation of 1000 was utilized. In IPA, the two transcriptionally responding animals at 3 dpi (M08 and M09) were used to identify signaling patterns at this timepoint, while all 3^rd^ trimester animals were analyzed at 7 dpi. Volcano plots, heatmaps, and Venn diagrams were generated using the EnhancedVolcano, pheatmap, and VennDiagram packages in R, respectively. For heatmaps of read count data, log2-transformed read counts of genes responsible for core enrichment of the indicated gene sets are plotted. Differential expression data from a previous cohort of male rhesus and cynomolgus macaques infected with the identical ZIKV isolate (Rio-U1) (Schouest et al., 2019, PREPRINT) is shown in Fig. 4. For this analysis, transcriptional profiles of immune signaling were generated using the nCounter NHP Immunology Panel of 770 macaque immune response genes (NanoString Technologies). RNA had been extracted from PAXgene blood RNA tubes (PreAnalytiX) using the PAXgene blood RNA kit (PreAnalytiX), and cDNA was synthesized using the RT2 First Strand Kit (Qiagen). Transcriptional responses were assessed at 3 dpi relative to expression levels pre-infection using nSolver software v4.0 (NanoString Technologies). Only DEGs that were present in both datasets (previous male cohort and 3^rd^ trimester animals at each timepoint) were included in comparison analyses.

### Anti-ZIKV binding antibody titers

Serum was tested for reactivity to ZIKV antigen using the commercial enzyme-linked immunospot assay (ELISA, Xpressbio) from before the initial infection, 28 days after the initial infection, on the day of reinfection, and 30 days after rechallenge. Responses during primary infection, reported as optical density (OD), were tested at a 1:50 dilution. This dilution proved too concentrated for the rechallenge, so a 1:200 dilution was used.

### Flow cytometry

Cryopreserved PBMCs were thawed and labeled with the following antibodies: CD16 AL488, CD169 PE, CD28 PE-CF594, CD95 PCP-Cy5.5, CD3 PE-Cy7, CD8 Pacific Blue, CD14 BV605, HLA-DR BV650, CD69 BV711, NKG2A APC, and CD4 APC-H7, followed by fixation, permeabilization, and labeling with an antibody against Ki67 AL700. Flow cytometry data were collected on a BD LSR II instrument and analyzed using FlowJo v10.

## Results

### CD8 lymphocyte depletion and rechallenge

CD8α lymphocyte depletion (targeting primarily CD8 T cells and NK cells) resulted in a rapid decrease in CD8+ cell counts to an undetectable level (Fig. 1a). Given that ZIKV can persist in tissues long past the clearance of virus from the serum, viral loads were determined prior to rechallenge to ensure CD8 depletion did not result in recrudescent viremia from a cryptic reservoir, and none was detected (data not shown). Following rechallenge, viral RNA was not detected by qRT-PCR in any sample at any timepoint, suggesting complete immunity to the Puerto Rican strain (Fig. 1b).

**Figure 1:**
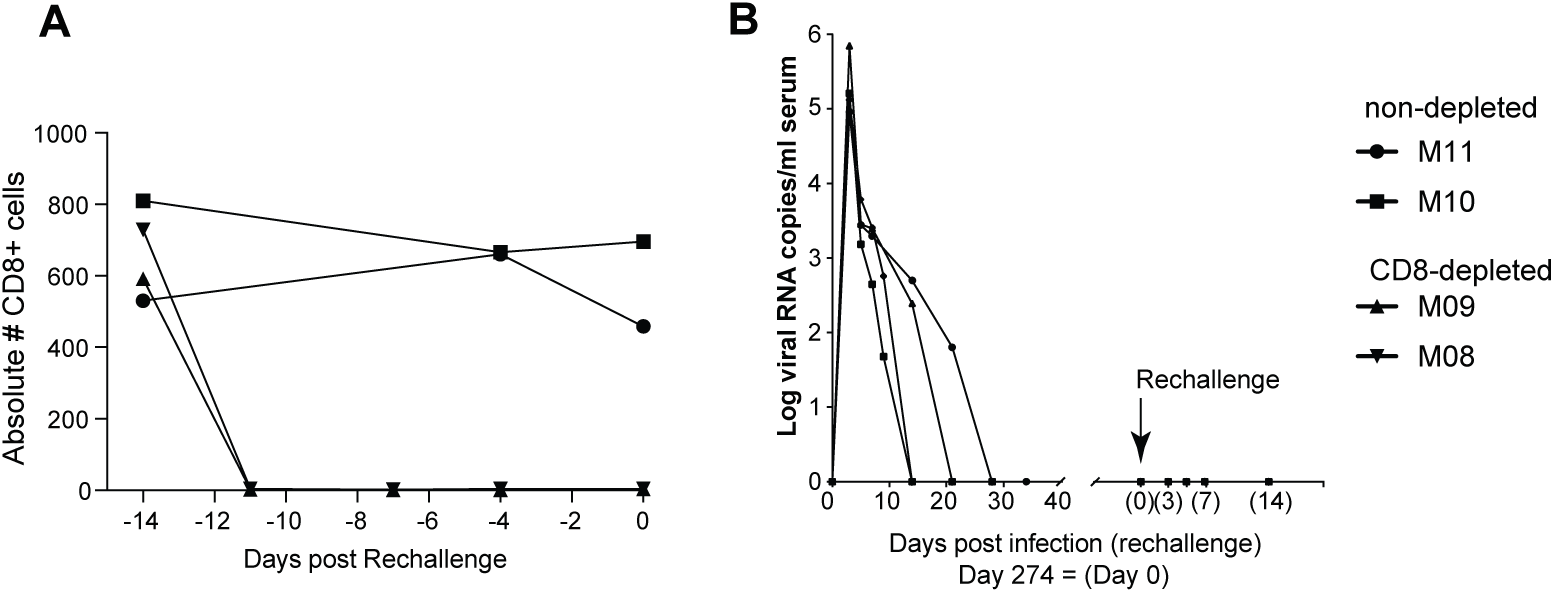
Study design. (**A**) CD8α+ cells were depleted in two of four 3^rd^ trimester animals using a 10-day protocol. Absolute counts of CD8+ cells in the blood dropped to near zero before ZIKV rechallenge. (**B**) Nine months after initial infection during pregnancy, four macaques, including the two CD8 depleted animals, were challenged subcutaneously with 10^4^ PFU of a Puerto Rican ZIKV strain (PRVABC-59), and viral loads monitored for one month.

### Transcriptome analysis following primary challenge

Although the primary infection appeared to confer complete protection in that all animals resisted rechallenge, we carried out transcriptome analysis to assess the quality of immune responses mounted during pregnancy. Primary infection resulted in the up- and downregulation of many genes at 3 and 7 dpi (Fig. 2a-b), but PCA revealed that early changes in gene expression at 3 dpi were driven principally by only 2 of 4 animals (Fig. 2c). By 7 dpi, however, all animals showed more uniform responses (Fig. 2c) that were characterized by the differential expression of a greater number of genes (Fig. 2d). To identify how changes in gene expression affected global signaling patterns during infection, we carried out an unsupervised, phenotype-independent analysis by way of GSVA (Hänzelmann et al., 2013). Interestingly, the two transcriptionally responsive animals at 3 dpi appeared to show trends mirroring those seen in all animals at 7 dpi (Fig. 2e). Enriched gene sets generally related to inflammation, innate immunity, viral replication, cell cycle arrest, and cell death, while downregulated gene sets involved hormone signaling, neurotransmitter release, and small molecule transport (Fig. 2e).

**Figure 2:**
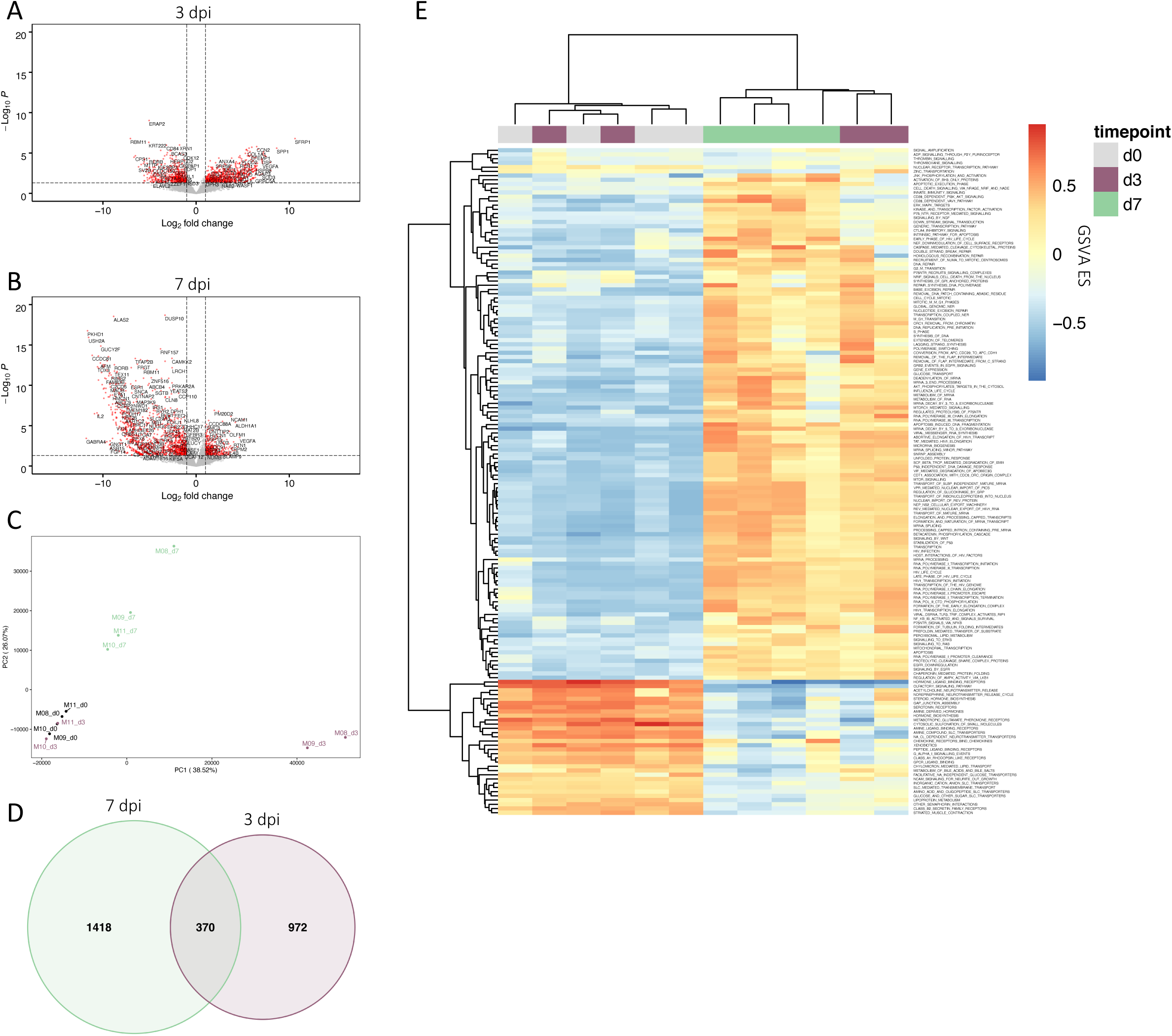
Transcriptome analysis of primary infection during pregnancy. (**A-D**) Volcano plots showing DEGs in PBMC at 3 dpi (A) and 7 dpi (B). (**C**) PCA plot showing the impacts of infection on the transcriptional landscape at 0, 3, and 7 dpi (d0, day 0; d3, day 3; d7, day 7; PC, principal component; consistent throughout). (**D**) Venn diagram showing the number of DEGs (p<0.05) at 3 and 7 dpi. (**E**) Heatmap showing GSVA values among significantly modulated gene sets. Gene sets in the Reactome databank were included if statistical significance was attained in pairwise comparisons among timepoints. (ES, enrichment score.)

Despite the up- and downmodulation of gene sets at 3 and 7 dpi (Fig. S1a-b) that in many cases appeared to be overlapping (Fig. 2e), a number of gene sets were differentially enriched at day 7 relative to day 3 (Fig. 3a), suggesting the possibility of divergent transcriptional signatures at these timepoints. Thus, we carried out a more detailed pathway analysis focusing on the progression of signaling patterns. Although there were fewer DEGs identified at 3 dpi relative to 7 dpi (Fig. 2d), a more varied functional fingerprint was evident at day 3 (Fig. 3b). GSEA of Reactome networks showed that maintenance of structural proteins was affected at either timepoint, though diseases associated with metabolism and protein modification were detected only at 3 dpi (Fig. 3b-c). GSEA additionally revealed some level of immune activation at 3 dpi, with an effect on neutrophil degranulation and platelet activation (Fig. S1c). Interestingly, GAS6 was upregulated at 3 dpi (Fig. S1c), which is a bridging molecule that facilitates binding of ZIKV virions to the putative entry receptor AXL (Meertens et al., 2017). Gamma-carboxylation was also induced at this timepoint (Fig. S1c), a function important in the binding of GAS6 to AXL (Geng et al., 2017).

**Figure 3:**
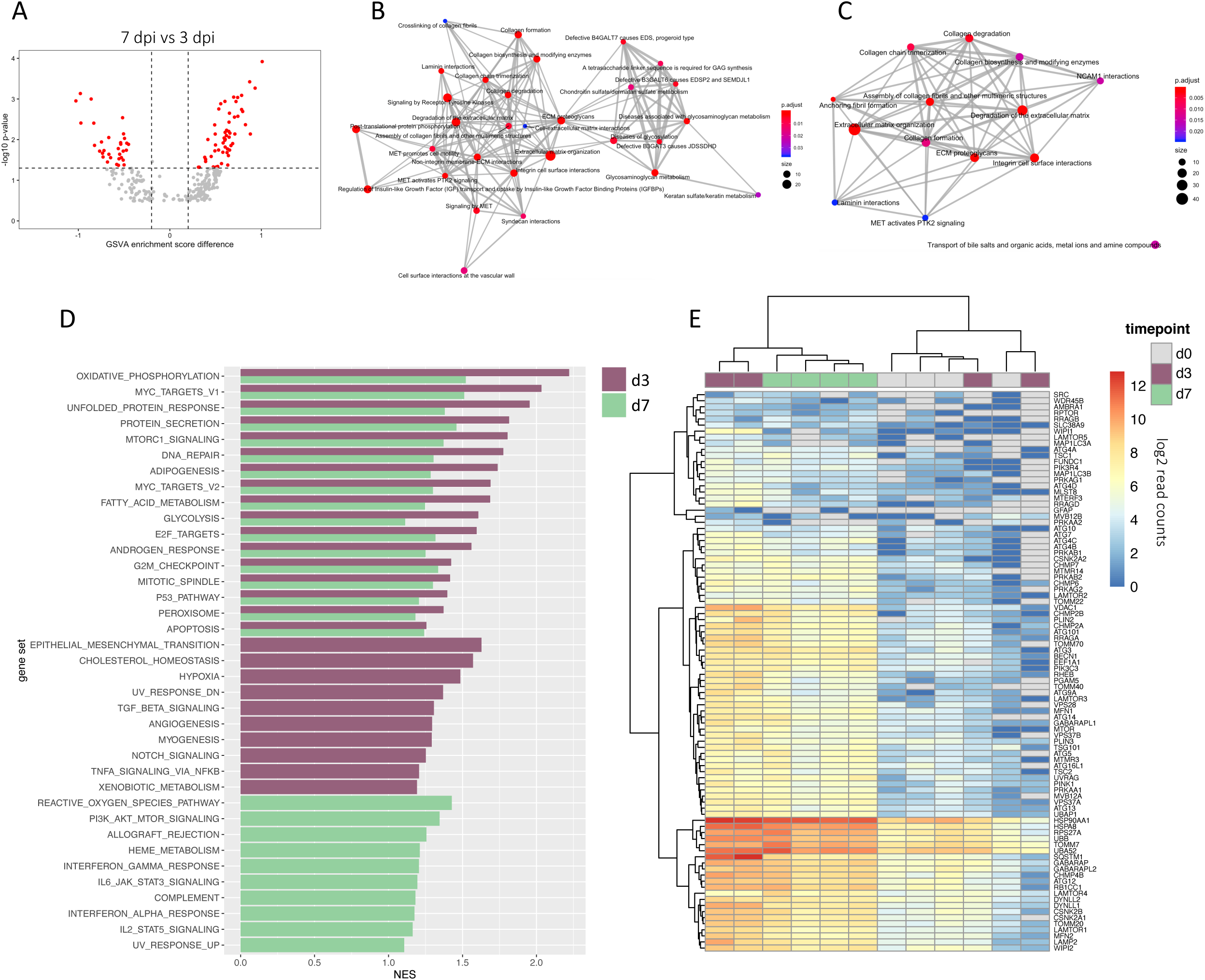
Transcriptional signatures of infection during pregnancy. (**A**) Volcano plot showing gene sets significantly modulated at 7 dpi relative to 3 dpi. (**B-C**) Enrichment map plots showing the interrelatedness of gene set networks at 3 dpi (B) and 7 dpi (C). (**D**) Gene sets from the Hallmarks collection that were significantly enriched (FDR<25%) by GSEA in 3^rd^ trimester animals at both timepoints (*top*) or only at 3 dpi (*middle*) or 7 dpi (*bottom*). (**E**) Heatmap showing read count data for autophagy related genes.

Viral infection is often associated with perturbations to metabolic processes, so to further characterize these effects which were initially identified by GSVA, we carried out targeted GSEA to compare phenotypes at 3 and 7 dpi to pre-infection. Indeed, we found that gene sets relating to cell respiration and lipid metabolism were among the most highly induced functions, and enrichment of these gene sets was generally greatest at 3 dpi (Fig. 3d). At day 3, there were additional signs of immunomodulation, including TGFβ signaling and angiogenesis, but by day 7, it was apparent that more of an inflammatory phenotype had emerged, marked by interferon (IFN) and proinflammatory cytokine signaling (Fig. 3d). The metabolic reprogramming at 3 dpi was characterized by changes in cell respiration (oxidative phosphorylation, glycolysis) and lipid metabolism (adipogenesis, fatty acid metabolism, cholesterol homeostasis, peroxisome) (Fig. 3d) that at the gene level showed activation in only the two early responding animals (Fig. S1d-e). Overarching effects on cell respiration and lipid metabolism might be explained by the induction of autophagy, which was also significantly enriched at 3 dpi (Fig. 3e). Autophagy has been shown to promote maternal-to-fetal transmission of ZIKV in mice through metabolic reprogramming in placental trophoblasts (Cao et al., 2017), and interestingly, one of the animals that showed early upregulation of autophagy signaling also had virus detectable in the amniotic fluid at multiple timepoints (Fig. S2a).

In contrast to the robust IFN responses that were evident in a previous cohort of male macaques infected with the identical ZIKV strain (Fig. 4a), the 3^rd^ trimester females showed comparatively muted innate immune responses at either timepoint. The female cohort showed limited evidence of inflammation (platelet degranulation and pattern recognition) that partially overlapped with signaling patterns previously observed in the males (Fig. 4b-c), but IPA rather pointed to an immunomodulatory phenotype in the 3^rd^ trimester animals at 3 dpi (Fig. S2b). In the males, transcriptional responses at day 3 indicated robust IFN signaling (Fig. 4a) and high levels of leukocyte homing and activation (Schouest et al., 2019, PREPRINT), while the same pathways were depressed in 3^rd^ trimester animals at the equivalent timepoint (Fig. S2b). Signs of immunoregulation were evident at 3 dpi, characterized by a lack of immune cell recruitment and activation that was driven by a decrease in a core set of chemokines (IL2) and their receptors (CCR7, IL12RB1), together with downregulated adhesion proteins (SELP, CD48, SELL, ICOS, CD40LG), signaling molecules (IRF1, NFATC2), and activation markers (CD69, CD48) (Fig. S2b). By 7 dpi, there was downregulation of estrogen receptor (ESR1) and other genes relating to fertility (AR, CCNE2) and organ development (CAV1, PPARGC1A) (Fig. S2c), implying a negative impact on reproductive function. IPA predicted the up- and downregulation of several immunomodulatory molecules in 3^rd^ trimester animals at both timepoints (Fig. S2d), including FGF2, which is known to support ZIKV infection by suppressing IFN signaling (Limonta et al., 2019). A number of other inflammatory pathways and immunomodulators such as TGFβ, IL10, and type-I IFN were also inversely regulated between 3 and 7 dpi (Fig. S2d-e), suggesting a complex regulation of immunity that was not present in non-pregnant animals infected with the same virus. We caution that transcriptional responses in the previous male cohort were profiled using NanoString technology, which is a more targeted platform compared to RNAseq, but the absence of robust antiviral signaling in the pregnant animals nonetheless suggests patterns of transcriptional activation that differed fundamentally from non-pregnant macaques.

**Figure 4:**
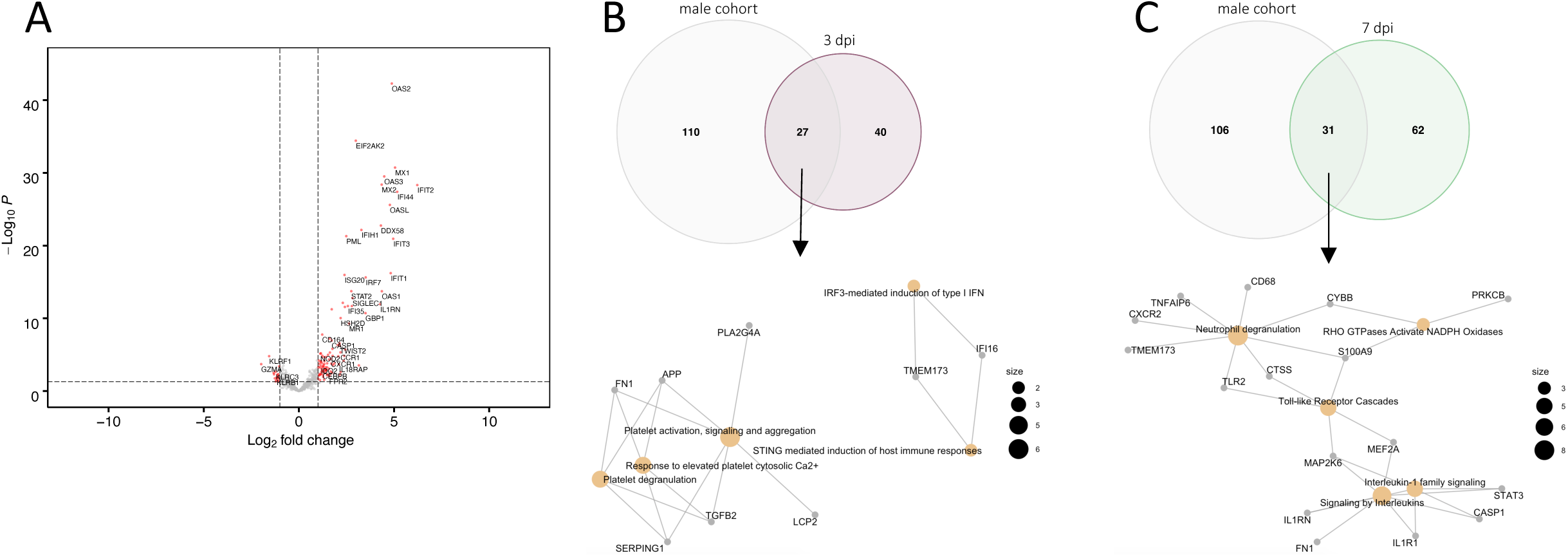
Comparison of gene expression patterns in pregnant animals to a previous male cohort. (**A**) Volcano plot showing DEGs from a previous cohort of male rhesus and cynomolgus macaques infected with ZIKV Rio-U1 (Schouest et al., 2019). DEGs were calculated from NanoString transcriptional analysis of immune responses in whole blood at 3 dpi relative to 0 dpi. (**B-C**) Comparison of DEGs among the previous male cohort and pregnant animals at 3 dpi (B) and 7 dpi (C). Pathway enrichment (*bottom*) was carried out on DEGs in common among the cohorts (*top*).

### Sustained anti-ZIKV antibody titers expand after rechallenge

Despite transcriptional evidence of an immunomodulatory phenotype during the initial challenge, binding antibodies were detected at 28 dpi using a 1:50 dilution of blood plasma, the only post-infection timepoint tested (Fig. 5a). Binding antibodies were re-assessed on the day of rechallenge, again using a 1:50 dilution, to provide a reference for determining humoral responses to a secondary infection. However, at this dilution, antibody responses were outside of the detection range (OD > 3.5) (data not shown), suggesting they had continued to rise since 28 dpi of the initial infection. We then repeated the assay using a 1:200 plasma dilution and found that the concentration of binding antibodies expanded after rechallenge in 3 of 4 animals, while antibodies in the fourth animal remained above the limit of detection (Fig. 5b).

**Figure 5:**
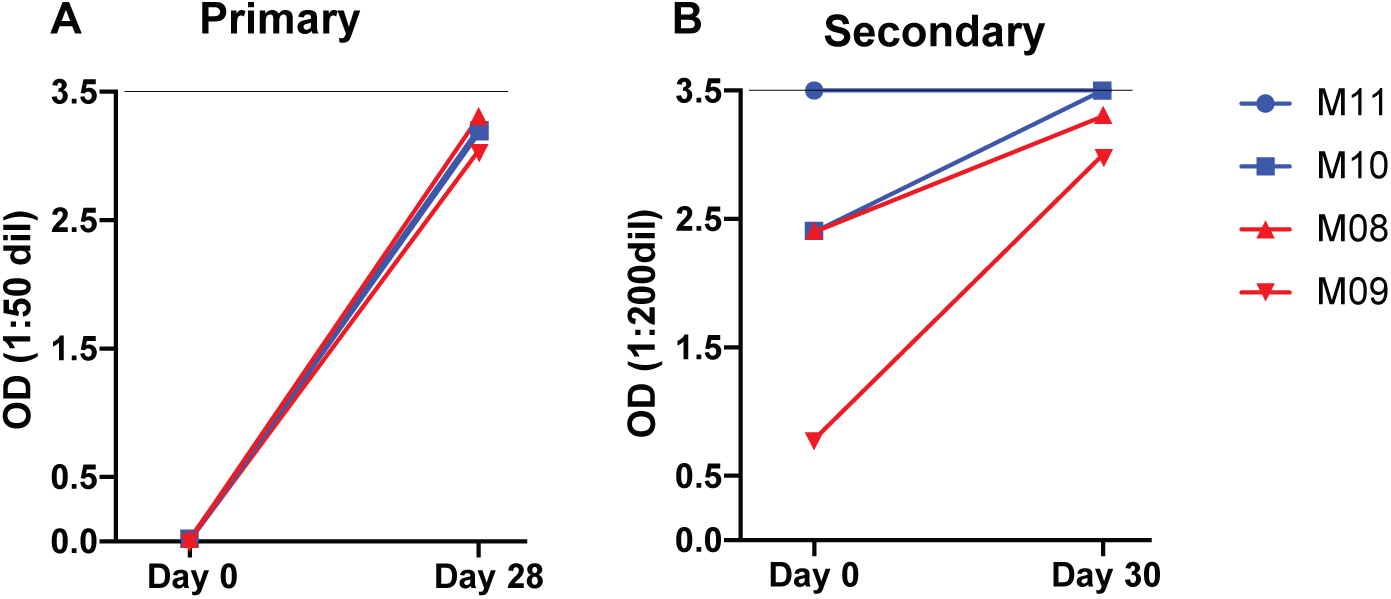
Anti-ZIKV humoral responses. (**A-B**) ELISA was used to assess anti-ZIKV humoral immunity after primary infection (A) and rechallenge (B). Antibody titers measured at a 1:50 dilution rose between the day of primary infection and day 28 (A). Anti-ZIKV antibody responses also expanded following rechallenge (B), which were tested at a 1:200 dilution as 1:50 proved too concentrated for the dynamic range of the assay.

### Immune activation following secondary challenge

We also assessed the activation of innate and adaptive immune cells as a surrogate of infection, given that viral RNA was not detected in the serum of any animal following rechallenge. Using a multicolor flow cytometry panel that we adapted from a previous ZIKV study (Schouest et al., 2019), we evaluated the proliferation (Ki67) and activation (CD69 or CD169) of T cells and monocyte subsets cells before and after rechallenge. CD169 is a biomarker of inflammation that has been used to track monocyte activation during acute ZIKV infection in macaques (Hirsch et al., 2017).

Classical and intermediate monocytes showed no discernable changes in frequency or activation following secondary challenge (Fig. 6a-f). However, nonclassical monocytes (CD14^-/low^, CD16^+^) expanded at 3 dpi in CD8-depleted animals (Fig. 6g). Although nonclassical monocytes showed no change in CD169 expression (Fig. 6h), there was an increase in activation as measured by CD69 expression predominantly in nondepleted animals at 5 dpi (Fig. 6i).

**Figure 6:**
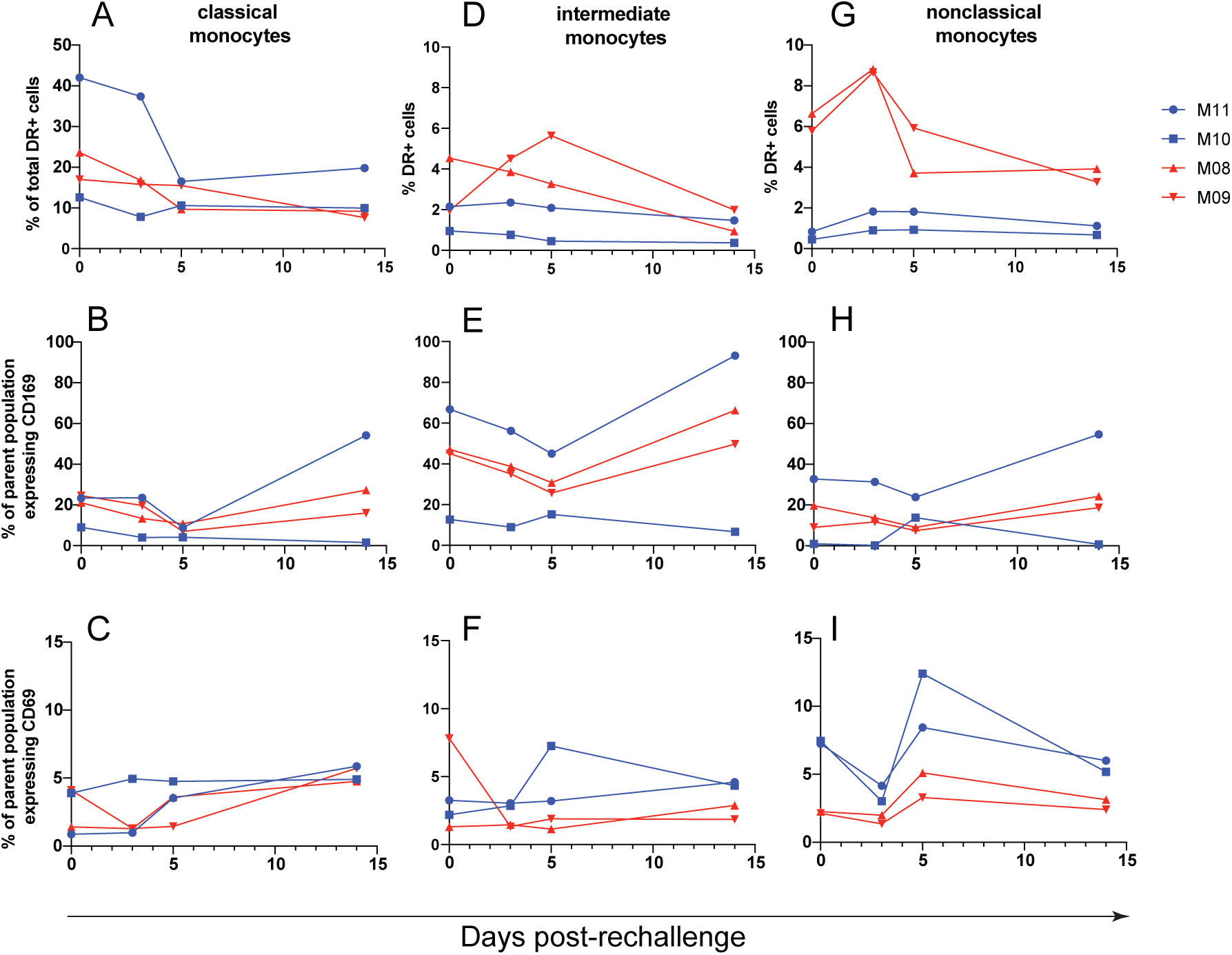
Monocyte changes after ZIKV rechallenge. Classical (CD14^+^, CD16^-^) (**A-C**), intermediate (CD14^+^, CD16^+^) (**D-F**), and non-classical (CD14^low/-^, CD16^+^) (**G-I**) monocytes were assessed for changes in frequency (A, D, G) and activation as measured by CD69 expression (B, E, H) and CD169 expression (C, F, I), after ZIKV rechallenge.

Following rechallenge, central memory CD4 T cells expanded in frequency primarily in CD8-depleted animals (Fig. 7a), and these cells also showed increases in activation (CD69 expression, Fig. 7b) and proliferation (Ki67 expression, Fig. 7c) between 3-5 dpi in the same animals. Nondepleted animals showed an increase in central memory CD4 T cell proliferation during the same period (Fig. 7c), but the magnitude of this increase was marginal compared to that of the CD8-depleted animals. Effector memory CD4 T cells showed a modest increase in frequency in CD8-depleted animals at 3-5 dpi (Fig. 7d) without clear changes in activation or proliferation (Fig. 7e-f). Naïve CD4 T cells showed a striking drop in frequency between 3-5 dpi, which was most pronounced in CD8-depleted animals (Fig. 7g). The decline in naïve CD4 T cell frequency was concomitant with an increase in activation (CD69 expression, Fig. 7h) and proliferation (Ki67 expression, Fig. 7i) primarily in 3 of 4 animals.

**Figure 7:**
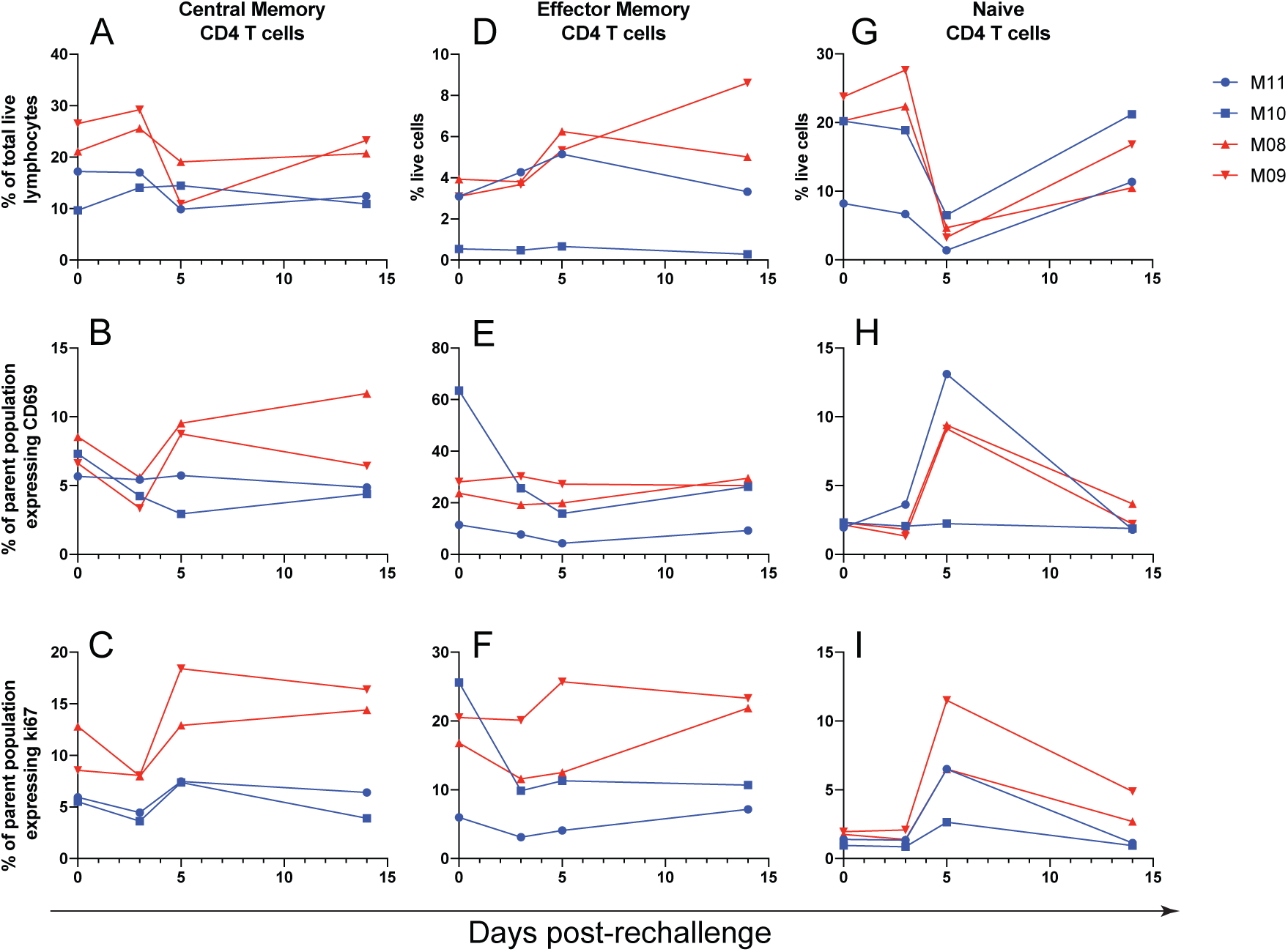
CD4 T cell changes after ZIKV rechallenge. Central memory (**A-C**), Effector memory (**D-F**), and naïve (**G-I**) CD4 T cells were assessed for changes in frequency (A, D, G), activation as measured by CD69 expression (B, E, H), and proliferation as measured by Ki67 expression (C, F, I), after ZIKV rechallenge.

Interestingly, phenotypic patterns in the CD8 T cell subsets of nondepleted animals generally mirrored those that occurred in the CD4 T cells of CD8-depleted animals. Although central memory CD8 T cells did not show appreciable changes in frequency (Fig. 8a) or activation (Fig. 8b) after rechallenge, there was a clear increase in proliferation between 3-5 dpi (Fig. 8c). Effector memory CD8 T cells expanded between the day of rechallenge and 5 dpi (Fig. 8d), but these cells did not become activated (Fig. 8e) and showed an increase in proliferation in only one of two nondepleted animals (Fig. 8f). In similar fashion to naïve CD4 T cells, naïve CD8 T cells dropped in frequency following rechallenge until 5 dpi (Fig. 8g) and showed marked increases in activation (Fig. 8h) and proliferation (Fig. 8i).

**Figure 8:**
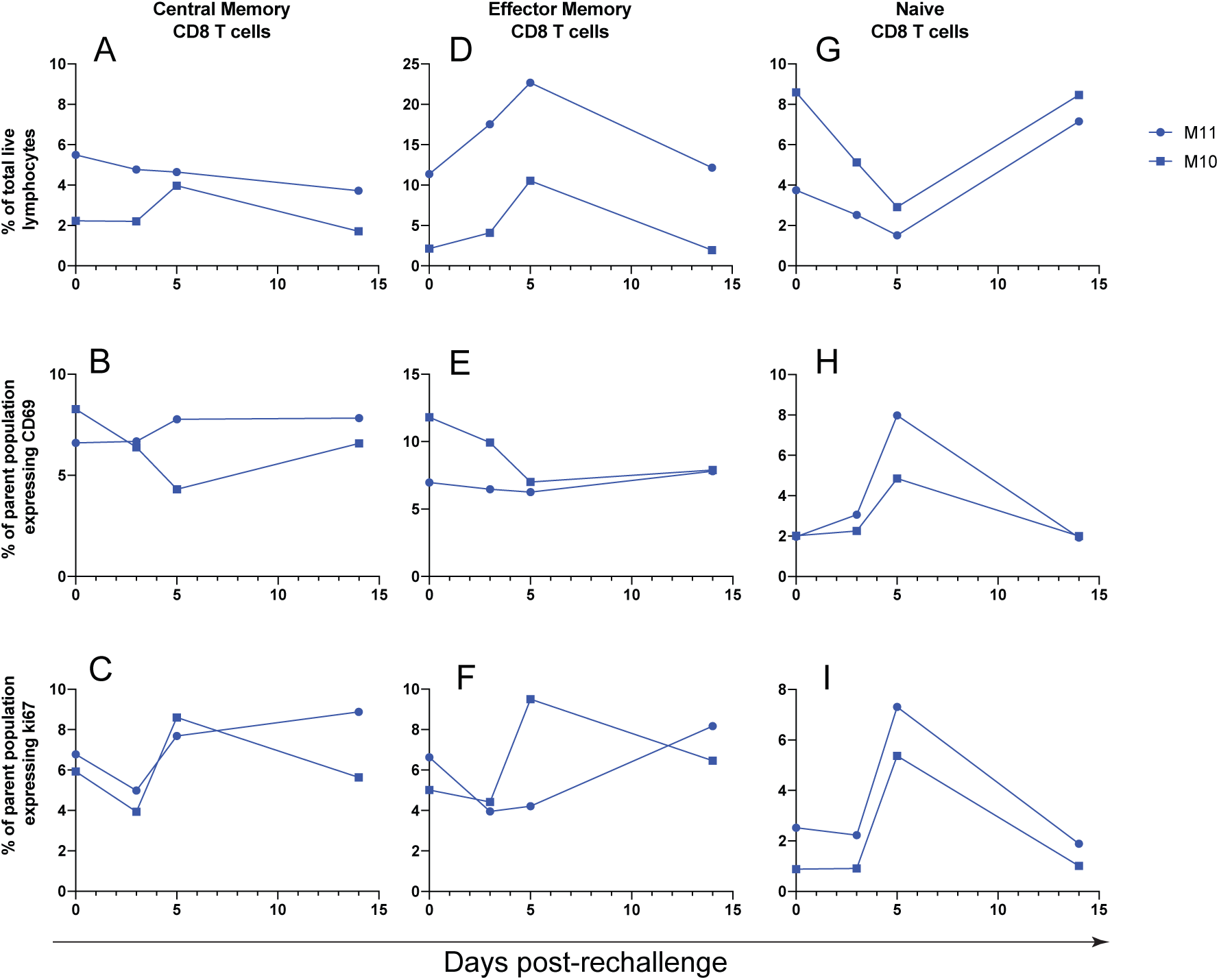
CD8 T cell changes after ZIKV rechallenge. Central memory (**A-C**), Effector memory (**D-F**), and naïve (**G-I**) CD8 T cells were assessed for changes in frequency (A, D, G), activation as measured by CD69 expression (B, E, H), and proliferation as measured by Ki67 expression (C, F, I), after ZIKV rechallenge.

## Discussion

ZIKV has been a known teratogen for some time, but the impacts of pregnancy on the quality of virus-specific immune responses have yet to be fully understood. The decrease in ZIKV incidence in the Western hemisphere since the peak of the outbreak in 2015 is of little reassurance until effective vaccines and therapeutics are mobilized. Although ZIKV vaccine candidates have performed well in preclinical settings, the congenital risks associated with ZIKV infection introduce a set of challenges to vaccine development that requires special consideration and carefully chosen animal models. Pregnancy presents a substantially altered immunologic state to facilitate fetal development and protect the mother and developing fetus from infectious agents (Kourtis et al., 2014), so it follows that immune correlates of protection during pregnancy might differ from mechanisms that are important in nonpregnant individuals.

Whether immune responses induced during pregnancy, due to either infection or vaccination, is sufficient to protect from subsequent infection has begun to be examined in murine and NHP models. A recent study in IFN-deficient mice showed that a live-attenuated vaccine protected pregnant dams from infection and also prevented *in utero* transmission, and this protection appeared to be mostly dependent on neutralizing antibodies (Shan et al., 2019). The report cautioned that higher antibody titers were required to protect pregnant animals compared to nonpregnant animals, and pregnancy appeared to negatively impact the potency of the T cell response induced by the vaccine, which are important considerations in the evaluation of future vaccine candidates. A separate study in NHPs showed that animals initially challenged during pregnancy mount immune responses similar to non-pregnant animals, and these responses adequately protect against a secondary challenge (Moreno et al., 2019). Again, antibodies appeared to be the most important correlate of protection since cell-mediated responses were not detected following reinfection. Our data similarly demonstrate that primary infection during pregnancy provides some level of protection; the presence of serum antibodies that continued to rise following rechallenge suggests a humoral response that might have limited viremia on secondary challenge. However, previous work has indicated that the best indicator of immunity to the related flavivirus dengue virus (DENV) in NHPs is not the absence of viremia but the lack of an anamnestic response following secondary challenge (Halstead et al., 1973). Although infection of NHPs with ZIKV typically produces a rapid serum viremia within days (Dudley et al., 2016; Magnani et al., 2018), the absence of virus following rechallenge in the present study does not exclude the possibility of low-level viral replication that we failed to detect. In either case, it is clear that immune responses mounted during pregnancy confer some level of protection to reinfection. Such a finding is not entirely surprising, as several studies have shown an efficient generation of immunity by vaccines administered during pregnancy (Healy, 2012; Munoz et al., 2014; Ohfuji et al., 2011; Sperling et al., 2012). Outdated models portray pregnancy as a global suppression of immunity (Mor et al., 2017), but these perspectives are no longer generally accepted, as it has become clear that pregnancy is rather a complex alteration of particular immune subsets to balance fetal development and protection from infection (Kourtis et al., 2014). Indeed, pregnancy is a progressive biological process that requires a progressively adapting immune microenvironment (Mor et al., 2017).

Our data show phenotypic changes in immune cell populations in both CD8-depleted and nondepleted animals, indicating that some level of cellular immune involvement might have conferred resistance to rechallenge. Nondepleted animals showed preferential expansion of effector memory CD8 T cells, while CD8-depleted animals showed greater increases in memory CD4 T cell subsets, possibly suggesting a compensatory CD4 response as we have observed previously in a cohort of male macaques that were similarly CD8 depleted prior to ZIKV challenge (Schouest et al., 2019). Antibody-mediated depletion experiments in mice have also illustrated the redundancy of adaptive responses to ZIKV, with the depletion of individual immune cell populations resulting in alternative compensatory responses (Scott et al., 2018). Together, these studies begin to reveal the plasticity of immune responses to ZIKV that may coordinate to maintain overall immune integrity.

Although limited sample availability precluded analysis of antigen specific CD8+ T cell responses, the potential for CD8+ lymphocytes to aid in protection from rechallenge is intriguing. Vaccine induced CD8 T cells are important in protection from Ebola virus challenge (Gupta et al., 2005; Sullivan et al., 2011; Warfield et al., 2005), and the lack of CD8 T cell epitopes in the currently licensed DENV vaccine (Dengvaxia, Sanofi Pasteur) might contribute to some of the efficacy concerns associated with vaccination (Tian et al., 2019). Whether this experimental readout will also be true for ZIKV immunity in humans is unknown, but these findings nonetheless underscore the importance of CD8 responses in protection from these viruses generally and in vaccine design. ZIKV-specific CD8 T cells are described in multiple species (Elong Ngono et al., 2017; Grifoni et al., 2018; Huang et al., 2017; Pardy et al., 2017) and may be important for viral clearance in mouse models. CD8 T cells have active roles in controlling infections caused by other flaviviruses including West Nile virus (Grifoni et al., 2018; Klein et al., 2005; Shrestha and Diamond, 2004; Shrestha et al., 2006; Wang et al., 2006; Wang et al., 2003), DENV (de Alwis et al., 2016; Lam et al., 2017; Regla-Nava et al., 2018; Rivino and Lim, 2017; Shi et al., 2015; Yauch et al., 2009), and yellow fever virus (Akondy et al., 2009; Bassi et al., 2015; Co et al., 2009; Nogueira et al., 2013), which, given their relatedness, implies that CD8 T cells may be similarly important in limiting ZIKV infection.

Phenotypic changes among monocyte subsets were minimal, but monocytes are the primary targets of ZIKV in the blood (Foo et al., 2017; Michlmayr et al., 2017; O’Connor et al., 2018), so any alterations in frequency or activation of these cells are potentially interesting. The increases in nonclassical monocyte frequency and activation in CD8-depleted and nondepleted animals are intriguing because among monocytes, the nonclassical subset is preferentially targeted by ZIKV infection (O’Connor et al., 2018). Moreover, in pregnant women, Asian-lineage ZIKV infection selectively expands nonclassical monocytes and induces an M2-skewed immunosuppressive phenotype (Foo et al., 2017). Why nonclassical monocytes responded differently among CD8-depleted and nondepleted animals is unclear, but similar patterns occurred in a previous study from our group that also used CD8 depletion in a cohort of male rhesus macaques (Schouest et al., 2019). In that study, the collateral depletion of NK cells in CD8-depleted animals appeared to skew patterns of monocyte activation among treatment groups, which might have also been the case here. The previous male cohort also showed strong upregulation of the activation marker CD169 on monocytes during acute infection (Schouest et al., 2019), but in contrast, the pregnant animals showed little fluctuation in CD169 expression, consistent with the lack of viremia in these animals. Together, these cellular immune data suggest some involvement of innate and adaptive cellular immune responses to the rechallenge virus, but the complete absence of viremia in both groups confirms that protection from rechallenge was robust.

Despite this protection, transcriptome analysis offered a more nuanced view of the impacts of pregnancy on immune activation during infection. Patterns of transcriptional activation in these animals differed fundamentally from non-pregnant macaques our group has previously inoculated with the same ZIKV strain, raising the possibility of immune modulation during pregnancy. For example, in a male cohort, we observed robust IFN signaling together with strong induction of immune cell activation and homing during acute infection (Schouest et al., 2019). While there was some evidence of inflammation and innate immunity in 3^rd^ trimester macaques, the transcriptional landscape in these animals was characterized by a level of complexity that also entailed metabolic and hormonal effects that might have ultimately skewed immune outcomes. The suppression of immune cell homing and activation was particularly intriguing and might imply an immunomodulatory phenotype, possibly owing to maternal hormonal regulation. Although pregnancy has, in the past, been viewed as an immunosuppressive host-graft relationship to prevent fetal damage, current models recognize pregnancy as a progression of stages, each requiring unique immunological cues (Mor et al., 2017). In the pregnant animals, immune activation was altered even among the two timepoints we obtained post-challenge, possibly reflecting a complex regulation of immunity to ZIKV infection in the context of pregnancy. Although we also detected several gene expression signatures that were predicted to negatively impact fetal development, any immunoregulatory phenotype that might have occurred ultimately did not compromise fetal health. Infants from two dams were born healthy and quickly cleared postnatal ZIKV challenge, and no adverse effects on nervous system development or behavior were noted, as described previously (Maness et al., 2019).

In addition to altered immune activation patterns, metabolic reprogramming through autophagy also appeared to characterize infection in these animals. Although autophagy generally aids in pathogen degradation and in the induction of immune responses during microbial infection (Ma et al., 2013), ZIKV and DENV, like other viruses, interact directly with autophagy pathways to promote their own replication (Chiramel and Best, 2018). Moreover, it has been shown in mice that ZIKV activates autophagy in placental trophoblasts to enhance vertical transmission (Cao et al., 2017), so it was fascinating that a pregnant macaque with evidence of early autophagy signaling patterns also had virus cross the placental barrier. Since autophagy is at its core a degradative process that frees biomolecules such as lipids to enter energy producing pathways, functions we detected in pregnant ZIKV infected animals, further metabolomic experiments should address whether autophagic flux is related to ZIKV infection generally or ZIKV infection during pregnancy specifically.

### Conclusions

Together, our data confirm findings from a recent study in NHPs suggesting that pregnancy does not overtly impair immune responses to ZIKV infection, a finding with potential implications for vaccine design. We caution that the cohort of animals used in immune phenotyping experiments was initially infected during the 3^rd^ trimester of pregnancy, so it remains possible that infection during the 1^st^ or 2^nd^ trimesters might produce less protective responses. Our findings add to a growing body of data describing the correlates of ZIKV-induced immunity in animal models of pregnancy, justifying vaccine research efforts in this unique subpopulation.

## Author contributions

**Blake Schouest**: methodology, formal analysis, investigation, writing – original draft, writing – review & editing, visualization. **Margaret H. Gilbert**: methodology, writing – review & editing, supervision. **Rudolf P Bohm**: writing – review & editing, supervision. **Faith Schiro**: investigation, writing – review & editing. **Pyone P. Aye**: project administration. **Antonito T Panganiban**: conceptualization, resources, writing – review & editing, funding acquisition. **Diogo Magnani**: conceptualization, methodology, writing – original draft, writing – review & editing. **Nicholas J Maness**: conceptualization, methodology, formal analysis, resources, writing – original draft, writing – review & editing, visualization, supervision, funding acquisition.

---

**Supplementary Figure 1:**
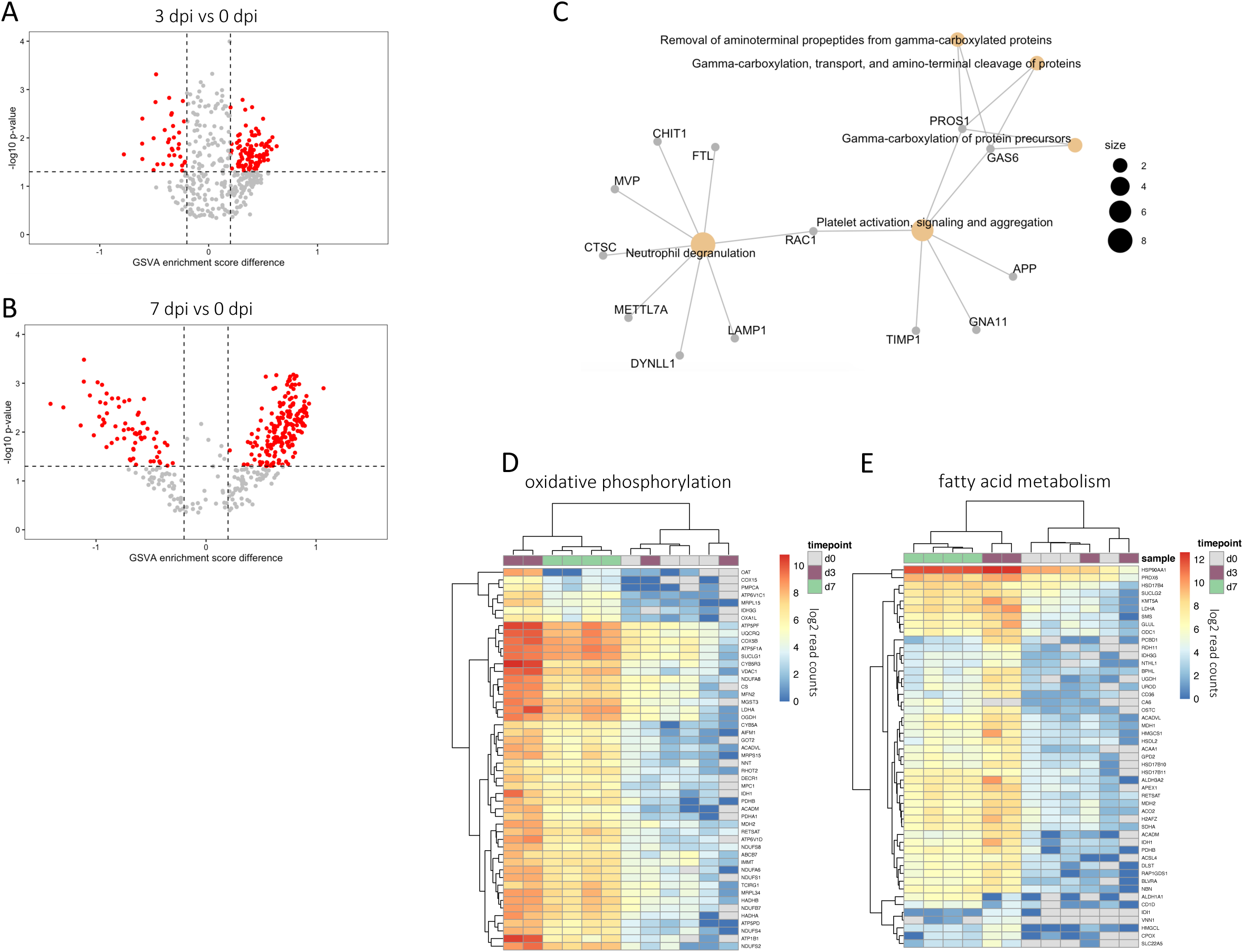
Pathway analysis of transcriptional signatures. (**A-B**) Volcano plots showing gene sets significantly modulated at 3 dpi (A) and 7 dpi (B) relative to pre-infection. (**C**) Gene network showing signaling patterns and genes induced at 3 dpi. (**D-E**) Heatmaps showing read count data for genes relating to oxidative phosphorylation (D) and fatty acid metabolism (E).

**Supplementary Figure 2:**
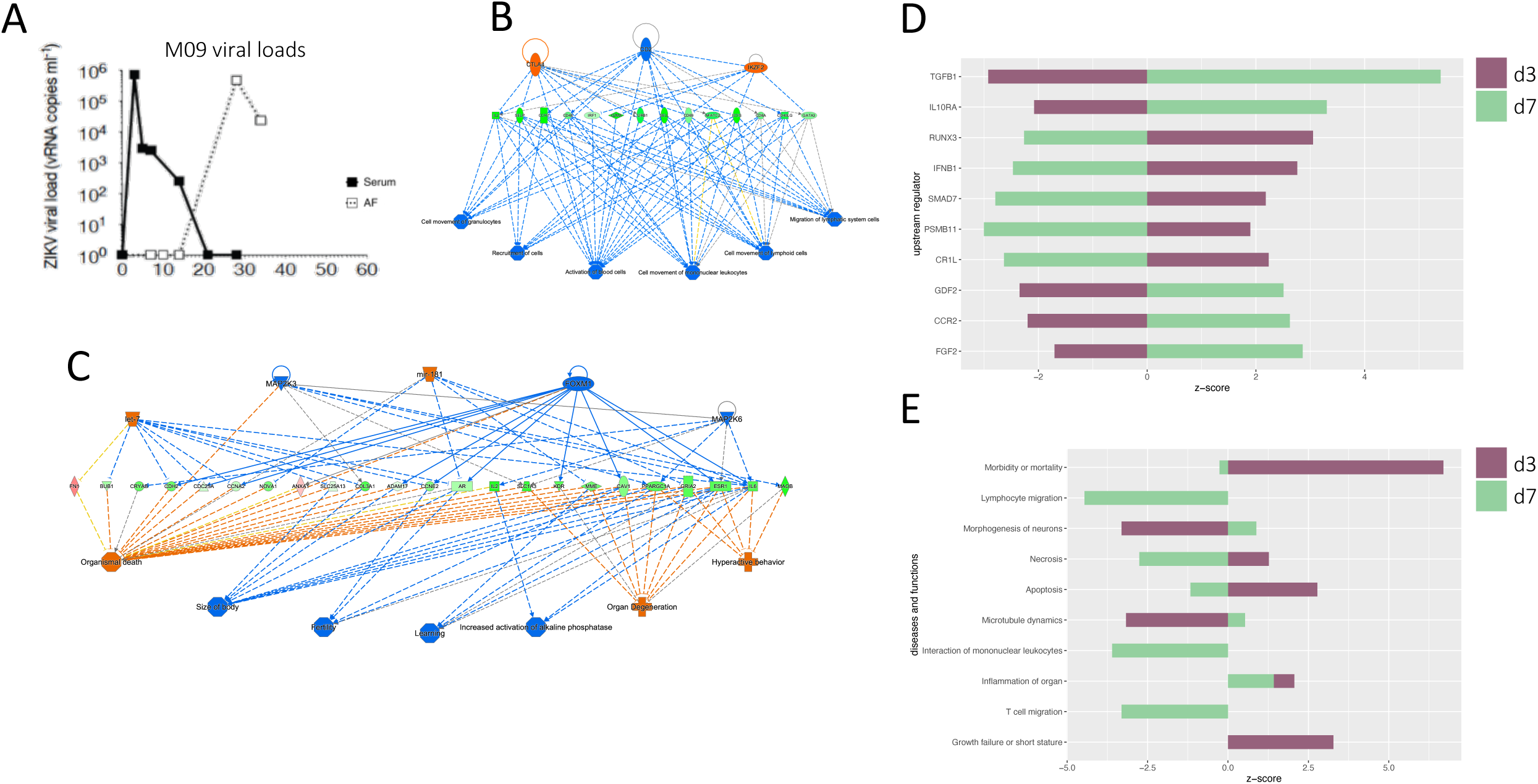
Putative consequences of gene expression patterns in pregnant animals. (**A**) Viral loads in serum and amniotic fluid (AF) during primary challenge in M09. (**B-C**) Regulator effects pathways from IPA, showing the predicted activation states of upstream regulators and canonical pathways at 3 dpi (B) and 7 dpi (C). (**D-E**) Predicted activation of biological regulators (E) and canonical pathways (F) at 3 and 7 dpi. Z-score represents predicted activation of the molecule or pathway.

## Bibliography

1 Akondy, R.S., Monson, N.D., Miller, J.D., Edupuganti, S., Teuwen, D., Wu, H., Quyyumi, F., Garg, S., Altman, J.D., Del Rio, C., Keyserling, H.L., Ploss, A., Rice, C.M., Orenstein, W.A., Mulligan, M.J., Ahmed, R., 2009. The yellow fever virus vaccine induces a broad and polyfunctional human memory CD8+ T cell response. J Immunol 183, 7919–7930.

2 Aliota, M.T., Dudley, D.M., Newman, C.M., Mohr, E.L., Gellerup, D.D., Breitbach, M.E., Buechler, C.R., Rasheed, M.N., Mohns, M.S., Weiler, A.M., Barry, G.L., Weisgrau, K.L., Eudailey, J.A., Rakasz, E.G., Vosler, L.J., Post, J., Capuano, S., 3rd, Golos, T.G., Permar, S.R., Osorio, J.E., Friedrich, T.C., O’Connor, S.L., O’Connor, D.H., 2016. Heterologous Protection against Asian Zika Virus Challenge in Rhesus Macaques. PLoS Negl Trop Dis 10, e0005168.

3 Araujo, L.M., Ferreira, M.L., Nascimento, O.J., 2016. Guillain-Barre syndrome associated with the Zika virus outbreak in Brazil. Arq Neuropsiquiatr 74, 253–255.

4 Avelino-Silva, V.I., Martin, J.N., 2016. Association between Guillain-Barre syndrome and Zika virus infection. Lancet 387, 2599.

5 Barton, M.A., Salvadori, M.I., 2016. Zika virus and microcephaly. CMAJ: Canadian Medical Association journal = journal de l’Association medicale canadienne 188, E118–119.

6 Bassi, M.R., Kongsgaard, M., Steffensen, M.A., Fenger, C., Rasmussen, M., Skjodt, K., Finsen, B., Stryhn, A., Buus, S., Christensen, J.P., Thomsen, A.R., 2015. CD8+ T cells complement antibodies in protecting against yellow fever virus. J Immunol 194, 1141–1153.

7 Bonaldo, M.C., Ribeiro, I.P., Lima, N.S., Dos Santos, A.A., Menezes, L.S., da Cruz, S.O., de Mello, I.S., Furtado, N.D., de Moura, E.E., Damasceno, L., da Silva, K.A., de Castro, M.G., Gerber, A.L., de Almeida, L.G., Lourenço-de-Oliveira, R., Vasconcelos, A.T., Brasil, P., 2016. Isolation of Infective Zika Virus from Urine and Saliva of Patients in Brazil. PLoS Negl Trop Dis 10, e0004816.

8 Brasil, P., Sequeira, P.C., Freitas, A.D., Zogbi, H.E., Calvet, G.A., de Souza, R.V., Siqueira, A.M., de Mendonca, M.C., Nogueira, R.M., de Filippis, A.M., Solomon, T., 2016. Guillain-Barre syndrome associated with Zika virus infection. Lancet 387, 1482.

9 Cao, B., Parnell, L.A., Diamond, M.S., Mysorekar, I.U., 2017. Inhibition of autophagy limits vertical transmission of Zika virus in pregnant mice. J Exp Med 214, 2303–2313.

10 Chiramel, A.I., Best, S.M., 2018. Role of autophagy in Zika virus infection and pathogenesis. Virus Res 254, 34–40.

11 Co, M.D., Kilpatrick, E.D., Rothman, A.L., 2009. Dynamics of the CD8 T-cell response following yellow fever virus 17D immunization. Immunology 128, e718–727.

12 Coffey, L.L., Pesavento, P.A., Keesler, R.I., Singapuri, A., Watanabe, J., Watanabe, R., Yee, J., Bliss-Moreau, E., Cruzen, C., Christe, K.L., Reader, J.R., von Morgenland, W., Gibbons, A.M., Allen, A.M., Linnen, J., Gao, K., Delwart, E., Simmons, G., Stone, M., Lanteri, M., Bakkour, S., Busch, M., Morrison, J., Van Rompay, K.K., 2017. Zika Virus Tissue and Blood Compartmentalization in Acute Infection of Rhesus Macaques. PloS one 12, e0171148.

13 de Alwis, R., Bangs, D.J., Angelo, M.A., Cerpas, C., Fernando, A., Sidney, J., Peters, B., Gresh, L., Balmaseda, A., de Silva, A.D., Harris, E., Sette, A., Weiskopf, D., 2016. Immunodominant Dengue Virus-Specific CD8+ T Cell Responses Are Associated with a Memory PD-1+ Phenotype. J Virol 90, 4771–4779.

14 Dick, G.W., Kitchen, S.F., Haddow, A.J., 1952. Zika virus. I. Isolations and serological specificity. Trans R Soc Trop Med Hyg 46, 509–520.

15 Dobin, A., Davis, C.A., Schlesinger, F., Drenkow, J., Zaleski, C., Jha, S., Batut, P., Chaisson, M., Gingeras, T.R., 2013. STAR: ultrafast universal RNA-seq aligner. Bioinformatics 29, 15–21.

16 Dudley, D.M., Aliota, M.T., Mohr, E.L., Weiler, A.M., Lehrer-Brey, G., Weisgrau, K.L., Mohns, M.S., Breitbach, M.E., Rasheed, M.N., Newman, C.M., Gellerup, D.D., Moncla, L.H., Post, J., Schultz-Darken, N., Schotzko, M.L., Hayes, J.M., Eudailey, J.A., Moody, M.A., Permar, S.R., O’Connor, S.L., Rakasz, E.G., Simmons, H.A., Capuano, S., Golos, T.G., Osorio, J.E., Friedrich, T.C., O’Connor, D.H., 2016. A rhesus macaque model of Asian-lineage Zika virus infection. Nature communications 7, 12204.

17 Ellul, M.A., Soares, C.N., Solomon, T., 2016. Zika virus and Guillain-Barre syndrome. J R Coll Physicians Edinb 46, 103–105.

18 Elong Ngono, A., Vizcarra, E.A., Tang, W.W., Sheets, N., Joo, Y., Kim, K., Gorman, M.J., Diamond, M.S., Shresta, S., 2017. Mapping and Role of the CD8(+) T Cell Response During Primary Zika Virus Infection in Mice. Cell host & microbe 21, 35–46.

19 Foo, S.S., Chen, W., Chan, Y., Bowman, J.W., Chang, L.C., Choi, Y., Yoo, J.S., Ge, J., Cheng, G., Bonnin, A., Nielsen-Saines, K., Brasil, P., Jung, J.U., 2017. Asian Zika virus strains target CD14+ blood monocytes and induce M2-skewed immunosuppression during pregnancy. Nat Microbiol 2, 1558–1570.

20 Geng, K., Kumar, S., Kimani, S.G., Kholodovych, V., Kasikara, C., Mizuno, K., Sandiford, O., Rameshwar, P., Kotenko, S.V., Birge, R.B., 2017. Requirement of Gamma-Carboxyglutamic Acid Modification and Phosphatidylserine Binding for the Activation of Tyro3, Axl, and Mertk Receptors by Growth Arrest-Specific 6. Front Immunol 8, 1521.

21 Grifoni, A., Costa-Ramos, P., Pham, J., Tian, Y., Rosales, S.L., Seumois, G., Sidney, J., de Silva, A.D., Premkumar, L., Collins, M.H., Stone, M., Norris, P.J., Romero, C.M.E., Durbin, A., Ricciardi, M.J., Ledgerwood, J.E., de Silva, A.M., Busch, M., Peters, B., Vijayanand, P., Harris, E., Falconar, A.K., Kallas, E., Weiskopf, D., Sette, A., 2018. Cutting Edge: Transcriptional Profiling Reveals Multifunctional and Cytotoxic Antiviral Responses of Zika Virus-Specific CD8. J Immunol 201, 3487–3491.

22 Grifoni, A., Pham, J., Sidney, J., O’Rourke, P.H., Paul, S., Peters, B., Martini, S.R., de Silva, A.D., Ricciardi, M.J., Magnani, D.M., Silveira, C.G.T., Maestri, A., Costa, P.R., de-Oliveira-Pinto, L.M., de Azeredo, E.L., Damasco, P.V., Phillips, E., Mallal, S., de Silva, A.M., Collins, M., Durbin, A., Diehl, S.A., Cerpas, C., Balmaseda, A., Kuan, G., Coloma, J., Harris, E., Crowe, J.E., Jr., Stone, M., Norris, P.J., Busch, M., Vivanco-Cid, H., Cox, J., Graham, B.S., Ledgerwood, J.E., Turtle, L., Solomon, T., Kallas, E.G., Watkins, D.I., Weiskopf, D., Sette, A., 2017. Prior Dengue virus exposure shapes T cell immunity to Zika virus in humans. J Virol.

23 Gupta, M., Greer, P., Mahanty, S., Shieh, W.J., Zaki, S.R., Ahmed, R., Rollin, P.E., 2005. CD8-mediated protection against Ebola virus infection is perforin dependent. J Immunol 174, 4198–4202.

24 Halstead, S.B., Casals, J., Shotwell, H., Palumbo, N., 1973. Studies on the immunization of monkeys against dengue. I. Protection derived from single and sequential virus infections. Am J Trop Med Hyg 22, 365–374.

25 Healy, C.M., 2012. Vaccines in pregnant women and research initiatives. Clin Obstet Gynecol 55, 474–486.

26 Hirsch, A.J., Smith, J.L., Haese, N.N., Broeckel, R.M., Parkins, C.J., Kreklywich, C., DeFilippis, V.R., Denton, M., Smith, P.P., Messer, W.B., Colgin, L.M., Ducore, R.M., Grigsby, P.L., Hennebold, J.D., Swanson, T., Legasse, A.W., Axthelm, M.K., MacAllister, R., Wiley, C.A., Nelson, J.A., Streblow, D.N., 2017. Zika Virus infection of rhesus macaques leads to viral persistence in multiple tissues. PLoS pathogens 13, e1006219.

27 Huang, H., Li, S., Zhang, Y., Han, X., Jia, B., Liu, H., Liu, D., Tan, S., Wang, Q., Bi, Y., Liu, W.J., Hou, B., Gao, G.F., Zhang, F., 2017. CD8+ T Cell Immune Response in Immunocompetent Mice during Zika Virus Infection. J Virol 91.

28 Hänzelmann, S., Castelo, R., Guinney, J., 2013. GSVA: gene set variation analysis for microarray and RNA-seq data. BMC Bioinformatics 14, 7.

29 Klein, R.S., Lin, E., Zhang, B., Luster, A.D., Tollett, J., Samuel, M.A., Engle, M., Diamond, M.S., 2005. Neuronal CXCL10 directs CD8+ T-cell recruitment and control of West Nile virus encephalitis. J Virol 79, 11457–11466.

30 Koide, F., Goebel, S., Snyder, B., Walters, K.B., Gast, A., Hagelin, K., Kalkeri, R., Rayner, J., 2016. Development of a Zika Virus Infection Model in Cynomolgus Macaques. Front Microbiol 7, 2028.

31 Kourtis, A.P., Read, J.S., Jamieson, D.J., 2014. Pregnancy and infection. N Engl J Med 370, 2211–2218.

32 Lam, J.H., Chua, Y.L., Lee, P.X., Martinez Gomez, J.M., Ooi, E.E., Alonso, S., 2017. Dengue vaccine-induced CD8+ T cell immunity confers protection in the context of enhancing, interfering maternal antibodies. JCI Insight 2.

33 Limonta, D., Jovel, J., Kumar, A., Lu, J., Hou, S., Airo, A.M., Lopez-Orozco, J., Wong, C.P., Saito, L., Branton, W., Wong, G.K., Mason, A., Power, C., Hobman, T.C., 2019. Fibroblast Growth Factor 2 Enhances Zika Virus Infection in Human Fetal Brain. J Infect Dis 220, 1377–1387.

34 Love, M.I., Huber, W., Anders, S., 2014. Moderated estimation of fold change and dispersion for RNA-seq data with DESeq2. Genome Biol 15, 550.

35 Ma, Y., Galluzzi, L., Zitvogel, L., Kroemer, G., 2013. Autophagy and cellular immune responses. Immunity 39, 211–227.

36 Magnani, D.M., Rogers, T.F., Maness, N.J., Grubaugh, N.D., Beutler, N., Bailey, V.K., Gonzalez-Nieto, L., Gutman, M.J., Pedreño-Lopez, N., Kwal, J.M., Ricciardi, M.J., Myers, T.A., Julander, J.G., Bohm, R.P., Gilbert, M.H., Schiro, F., Aye, P.P., Blair, R.V., Martins, M.A., Falkenstein, K.P., Kaur, A., Curry, C.L., Kallas, E.G., Desrosiers, R.C., Goldschmidt-Clermont, P.J., Whitehead, S.S., Andersen, K.G., Bonaldo, M.C., Lackner, A.A., Panganiban, A.T., Burton, D.R., Watkins, D.I., 2018. Fetal demise and failed antibody therapy during Zika virus infection of pregnant macaques. Nat Commun 9, 1624.

37 Maness, N.J., Schouest, B., Singapuri, A., Dennis, M., Gilbert, M.H., Bohm, R.P., Schiro, F., Aye, P.P., Baker, K., Van Rompay, K.K.A., Lackner, A.A., Bonaldo, M.C., Blair, R.V., Permar, S.R., Coffey, L.L., Panganiban, A.T., Magnani, D., 2019. Postnatal Zika virus infection of nonhuman primate infants born to mothers infected with homologous Brazilian Zika virus. Sci Rep 9, 12802.

38 Meertens, L., Labeau, A., Dejarnac, O., Cipriani, S., Sinigaglia, L., Bonnet-Madin, L., Le Charpentier, T., Hafirassou, M.L., Zamborlini, A., Cao-Lormeau, V.M., Coulpier, M., Missé, D., Jouvenet, N., Tabibiazar, R., Gressens, P., Schwartz, O., Amara, A., 2017. Axl Mediates ZIKA Virus Entry in Human Glial Cells and Modulates Innate Immune Responses. Cell Rep 18, 324–333.

39 Melo, A.S., Aguiar, R.S., Amorim, M.M., Arruda, M.B., Melo, F.O., Ribeiro, S.T., Batista, A.G., Ferreira, T., Dos Santos, M.P., Sampaio, V.V., Moura, S.R., Rabello, L.P., Gonzaga, C.E., Malinger, G., Ximenes, R., de Oliveira-Szejnfeld, P.S., Tovar-Moll, F., Chimelli, L., Silveira, P.P., Delvechio, R., Higa, L., Campanati, L., Nogueira, R.M., Filippis, A.M., Szejnfeld, J., Voloch, C.M., Ferreira, O.C., Jr., Brindeiro, R.M., Tanuri, A., 2016. Congenital Zika Virus Infection: Beyond Neonatal Microcephaly. JAMA neurology 73, 1407–1416.

40 Michlmayr, D., Andrade, P., Gonzalez, K., Balmaseda, A., Harris, E., 2017. CD14+CD16+ monocytes are the main target of Zika virus infection in peripheral blood mononuclear cells in a paediatric study in Nicaragua. Nat Microbiol 2, 1462–1470.

41 Moore, C.A., Staples, J.E., Dobyns, W.B., Pessoa, A., Ventura, C.V., Fonseca, E.B., Ribeiro, E.M., Ventura, L.O., Neto, N.N., Arena, J.F., Rasmussen, S.A., 2017. Characterizing the Pattern of Anomalies in Congenital Zika Syndrome for Pediatric Clinicians. JAMA Pediatr 171, 288–295.

42 Mor, G., Aldo, P., Alvero, A.B., 2017. The unique immunological and microbial aspects of pregnancy. Nat Rev Immunol 17, 469–482.

43 Moreno, G.K., Newman, C.M., Koenig, M.R., Mohns, M.S., Weiler, A.M., Rybarczyk, S., Weisgrau, K.L., Vosler, L.J., Pomplun, N., Schultz-Darken, N., Rakasz, E., Dudley, D.M., Friedrich, T.C., O’Connor, D.H., 2019. Long-term protection of rhesus macaques from Zika virus reinfection. J Virol.

44 Morrison, T.E., Diamond, M.S., 2017. Animal Models of Zika Virus Infection, Pathogenesis, and Immunity. J Virol 91.

45 Munoz, F.M., Bond, N.H., Maccato, M., Pinell, P., Hammill, H.A., Swamy, G.K., Walter, E.B., Jackson, L.A., Englund, J.A., Edwards, M.S., Healy, C.M., Petrie, C.R., Ferreira, J., Goll, J.B., Baker, C.J., 2014. Safety and immunogenicity of tetanus diphtheria and acellular pertussis (Tdap) immunization during pregnancy in mothers and infants: a randomized clinical trial. JAMA 311, 1760–1769.

46 Nguyen, S.M., Antony, K.M., Dudley, D.M., Kohn, S., Simmons, H.A., Wolfe, B., Salamat, M.S., Teixeira, L.B.C., Wiepz, G.J., Thoong, T.H., Aliota, M.T., Weiler, A.M., Barry, G.L., Weisgrau, K.L., Vosler, L.J., Mohns, M.S., Breitbach, M.E., Stewart, L.M., Rasheed, M.N., Newman, C.M., Graham, M.E., Wieben, O.E., Turski, P.A., Johnson, K.M., Post, J., Hayes, J.M., Schultz-Darken, N., Schotzko, M.L., Eudailey, J.A., Permar, S.R., Rakasz, E.G., Mohr, E.L., Capuano, S., 3rd, Tarantal, A.F., Osorio, J.E., O’Connor, S.L., Friedrich, T.C., O’Connor, D.H., Golos, T.G., 2017. Highly efficient maternal-fetal Zika virus transmission in pregnant rhesus macaques. PLoS pathogens 13, e1006378.

47 Nogueira, R.T., Nogueira, A.R., Pereira, M.C.S., Rodrigues, M.M., Neves, P., Galler, R., Bonaldo, M.C., 2013. Correction: Recombinant Yellow Fever Viruses Elicit CD8+ T Cell Responses and Protective Immunity against Trypanosoma cruzi. PloS one 8.

48 O’Connor, M.A., Tisoncik-Go, J., Lewis, T.B., Miller, C.J., Bratt, D., Moats, C.R., Edlefsen, P.T., Smedley, J., Klatt, N.R., Gale, M., Jr., Fuller, D.H., 2018. Early cellular innate immune responses drive Zika viral persistence and tissue tropism in pigtail macaques. Nature communications 9, 3371.

49 Ohfuji, S., Fukushima, W., Deguchi, M., Kawabata, K., Yoshida, H., Hatayama, H., Maeda, A., Hirota, Y., 2011. Immunogenicity of a monovalent 2009 influenza A (H1N1) vaccine among pregnant women: lowered antibody response by prior seasonal vaccination. The Journal of infectious diseases 203, 1301–1308.

50 Osuna, C.E., Lim, S.Y., Deleage, C., Griffin, B.D., Stein, D., Schroeder, L.T., Omange, R., Best, K., Luo, M., Hraber, P.T., Andersen-Elyard, H., Ojeda, E.F., Huang, S., Vanlandingham, D.L., Higgs, S., Perelson, A.S., Estes, J.D., Safronetz, D., Lewis, M.G., Whitney, J.B., 2016. Zika viral dynamics and shedding in rhesus and cynomolgus macaques. Nature medicine 22, 1448–1455.

51 Pardy, R.D., Rajah, M.M., Condotta, S.A., Taylor, N.G., Sagan, S.M., Richer, M.J., 2017. Analysis of the T Cell Response to Zika Virus and Identification of a Novel CD8+ T Cell Epitope in Immunocompetent Mice. PLoS pathogens 13, e1006184.

52 Plourde, A.R., Bloch, E.M., 2016. A Literature Review of Zika Virus. Emerging infectious diseases 22, 1185–1192.

53 Regla-Nava, J.A., Elong Ngono, A., Viramontes, K.M., Huynh, A.T., Wang, Y.T., Nguyen, A.T., Salgado, R., Mamidi, A., Kim, K., Diamond, M.S., Shresta, S., 2018. Cross-reactive Dengue virus-specific CD8(+) T cells protect against Zika virus during pregnancy. Nature communications 9, 3042.

54 Ricciardi, M.J., Magnani, D.M., Grifoni, A., Kwon, Y.C., Gutman, M.J., Grubaugh, N.D., Gangavarapu, K., Sharkey, M., Silveira, C.G.T., Bailey, V.K., Pedreno-Lopez, N., Gonzalez-Nieto, L., Maxwell, H.S., Domingues, A., Martins, M.A., Pham, J., Weiskopf, D., Altman, J., Kallas, E.G., Andersen, K.G., Stevenson, M., Lichtenberger, P., Choe, H., Whitehead, S.S., Sette, A., Watkins, D.I., 2017. Ontogeny of the B- and T-cell response in a primary Zika virus infection of a dengue-naive individual during the 2016 outbreak in Miami, FL. PLoS Negl Trop Dis 11, e0006000.

55 Ritchie, M.E., Phipson, B., Wu, D., Hu, Y., Law, C.W., Shi, W., Smyth, G.K., 2015. limma powers differential expression analyses for RNA-sequencing and microarray studies. Nucleic Acids Res 43, e47.

56 Rivino, L., Lim, M.Q., 2017. CD4(+) and CD8(+) T-cell immunity to Dengue - lessons for the study of Zika virus. Immunology 150, 146–154.

57 Schouest, B., Fahlberg, M., Scheef, E.A., Ward, M.J., Headrick, K., Szeltner, D.M., Blair, R.V., Gilbert, M.H., Doyle-Meyers, L.A., Danner, V.W., Bonaldo, M.C., Wesson, D.M., Panganiban, A.T., Maness, N.J., 2019. CD8+ lymphocytes modulate Zika virus dynamics and tissue dissemination and orchestrate antiviral immunity. bioRxiv 475418.

58 Scott, J.M., Lebratti, T.J., Richner, J.M., Jiang, X., Fernandez, E., Zhao, H., Fremont, D.H., Diamond, M.S., Shin, H., 2018. Cellular and Humoral Immunity Protect against Vaginal Zika Virus Infection in Mice. J Virol 92.

59 Shan, C., Xie, X., Luo, H., Muruato, A.E., Liu, Y., Wakamiya, M., La, J.H., Chung, J.M., Weaver, S.C., Wang, T., Shi, P.Y., 2019. Maternal vaccination and protective immunity against Zika virus vertical transmission. Nat Commun 10, 5677.

60 Shi, J., Sun, J., Wu, M., Hu, N., Li, J., Li, Y., Wang, H., Hu, Y., 2015. Inferring Protective CD8+ T-Cell Epitopes for NS5 Protein of Four Serotypes of Dengue Virus Chinese Isolates Based on HLA-A, -B and -C Allelic Distribution: Implications for Epitope-Based Universal Vaccine Design. PloS one 10, e0138729.

61 Shrestha, B., Diamond, M.S., 2004. Role of CD8+ T cells in control of West Nile virus infection. J Virol 78, 8312–8321.

62 Shrestha, B., Samuel, M.A., Diamond, M.S., 2006. CD8+ T cells require perforin to clear West Nile virus from infected neurons. J Virol 80, 119–129.

63 Sperling, R.S., Engel, S.M., Wallenstein, S., Kraus, T.A., Garrido, J., Singh, T., Kellerman, L., Moran, T.M., 2012. Immunogenicity of trivalent inactivated influenza vaccination received during pregnancy or postpartum. Obstetrics and gynecology 119, 631–639.

64 Subramanian, A., Tamayo, P., Mootha, V.K., Mukherjee, S., Ebert, B.L., Gillette, M.A., Paulovich, A., Pomeroy, S.L., Golub, T.R., Lander, E.S., Mesirov, J.P., 2005. Gene set enrichment analysis: a knowledge-based approach for interpreting genome-wide expression profiles. Proc Natl Acad Sci U S A 102, 15545–15550.

65 Sullivan, N.J., Hensley, L., Asiedu, C., Geisbert, T.W., Stanley, D., Johnson, J., Honko, A., Olinger, G., Bailey, M., Geisbert, J.B., Reimann, K.A., Bao, S., Rao, S., Roederer, M., Jahrling, P.B., Koup, R.A., Nabel, G.J., 2011. CD8+ cellular immunity mediates rAd5 vaccine protection against Ebola virus infection of nonhuman primates. Nature medicine 17, 1128–1131.

66 Tian, Y., Grifoni, A., Sette, A., Weiskopf, D., 2019. Human T Cell Response to Dengue Virus Infection. Front Immunol 10, 2125.

67 Wang, Y., Lobigs, M., Lee, E., Koskinen, A., Mullbacher, A., 2006. CD8(+) T cell-mediated immune responses in West Nile virus (Sarafend strain) encephalitis are independent of gamma interferon. J Gen Virol 87, 3599–3609.

68 Wang, Y., Lobigs, M., Lee, E., Mullbacher, A., 2003. CD8+ T cells mediate recovery and immunopathology in West Nile virus encephalitis. J Virol 77, 13323–13334.

69 Warfield, K.L., Olinger, G., Deal, E.M., Swenson, D.L., Bailey, M., Negley, D.L., Hart, M.K., Bavari, S., 2005. Induction of humoral and CD8+ T cell responses are required for protection against lethal Ebola virus infection. J Immunol 175, 1184–1191.

70 Xu, X., Vaughan, K., Weiskopf, D., Grifoni, A., Diamond, M.S., Sette, A., Peters, B., 2016. Identifying Candidate Targets of Immune Responses in Zika Virus Based on Homology to Epitopes in Other Flavivirus Species. PLoS Curr 8.

71 Yauch, L.E., Zellweger, R.M., Kotturi, M.F., Qutubuddin, A., Sidney, J., Peters, B., Prestwood, T.R., Sette, A., Shresta, S., 2009. A protective role for dengue virus-specific CD8+ T cells. J Immunol 182, 4865–4873.

72 Yu, G., He, Q.Y., 2016. ReactomePA: an R/Bioconductor package for reactome pathway 708 analysis and visualization. Mol Biosyst 12, 477–479.

73 Zanluca, C., Melo, V.C., Mosimann, A.L., Santos, G.I., Santos, C.N., Luz, K., 2015. First report of autochthonous transmission of Zika virus in Brazil. Memorias do Instituto Oswaldo Cruz 110, 711 569–572.

